# LiverHomo: An Integrated and Interactive Human Liver Single-Cell Atlas with Protein Interaction and Cell–Cell Communication Networks

**DOI:** 10.64898/2026.10.20.755227

**Authors:** Steven Yu, Ziyu Shi, Zhiyuan Song, Esteban Arroyave Sierra, Omar A. Saldarriaga, Heather L. Stevenson, Dongyin Guan, Bingning Dong, Haiqing Zhao

## Abstract

LiverHomo is an integrated single-cell atlas comprising 1,157,134 cells from 16 publicly available human single-cell RNA-sequencing (scRNA-seq) datasets, spanning healthy and fetal liver tissues, seven major liver diseases, and two tumor-associated contexts: metastatic lymph nodes and portal vein tumor thrombus. Building on existing liver single-cell resources, LiverHomo extends transcriptomic exploration by incorporating predicted protein–protein interaction (PPI) and cell–cell communication (CCI) networks. The resource provides an interactive web portal (https://www.liverhomo.org) for exploring cell-type distributions, gene expression patterns, differential expression, Gene Ontology annotations, and context-specific molecular and intercellular interaction networks across diseases and cell types. To make these multidimensional data easy and intuitive to access, the portal incorporates *Ask LiverHomo*, an AI-powered virtual assistant that interprets natural-language queries, including multilingual input, and retrieves atlas-derived results and visualizations through predefined analytical functions. Interactive expression plots, network visualizations, and cell-level metadata exploration let users investigate liver biology across cellular, transcriptomic, and molecular interaction levels. The underlying datasets and analytical outputs are available for download to support independent analysis and reuse. Together, LiverHomo provides an integrated and accessible resource for investigating cellular and molecular alterations across human liver health and disease.

## 1. Introduction

The liver is a highly heterogeneous organ that coordinates metabolic homeostasis, detoxification, and immune regulation through complex interactions among hepatocytes, cholangiocytes, endothelial cells, hepatic stellate cells, and diverse resident and infiltrating immune populations (1,2). Disruption and remodeling of these cellular populations and their interactions contribute to a broad spectrum of liver diseases, including metabolic dysfunction-associated steatotic liver disease (MASLD) or metabolic dysfunction-associated steatohepatitis (MASH), alcohol-associated liver disease (ALD), cholestatic diseases such as primary biliary cholangitis (PBC) and primary sclerosing cholangitis (PSC), fibrosis and cirrhosis, and primary liver cancers (3,4). Primary liver cancer represents a major global health burden and comprises two major histological forms, hepatocellular carcinoma (HCC) and intrahepatic cholangiocarcinoma (iCCA) (5,6). HCC typically develops in the setting of chronic hepatic injury, inflammation, fibrosis, and cirrhosis, whereas iCCA is associated with distinct pathological conditions, including cholangiopathies such as PSC, while sharing some metabolic risk factors with HCC (7–9). Their distinct pathological contexts also shape their cellular microenvironments: HCC frequently develops in chronically injured or cirrhotic, hepatocyte-rich parenchyma, whereas iCCA is characterized by a prominent desmoplastic and fibroblast-rich stroma (10,11). Understanding this diverse spectrum of liver diseases therefore requires resolving not only disease-associated molecular alterations but also the specific cellular contexts and intercellular interactions in which they occur.

Single-cell RNA sequencing (scRNA-seq) has substantially expanded our ability to resolve the human liver’s cellular complexity of the human liver and characterize disease-associated changes at cell-type resolution (12). Recent studies have revealed hepatocyte zonation, heterogeneous immune populations, disease-associated macrophage and fibroblast states, and transcriptional remodeling of epithelial and stromal populations across liver diseases (13). The rapidly growing collection of human liver scRNA-seq datasets therefore provides an opportunity to systematically investigate cellular and transcriptional alterations across diverse pathological conditions. However, researchers have largely generated these datasets independently across different patient cohorts, disease contexts, sequencing platforms, and analytical pipelines. Differences in quality control, cell annotation, and data processing further complicate direct comparisons across studies, limiting systematic investigation of shared and disease-specific cellular programs.

Several single-cell atlases and databases have been developed to improve the accessibility and integration of human liver transcriptomic data (14,15). Nevertheless, existing resources primarily focus on cellular composition and gene expression, whereas cellular phenotypes also emerge through molecular interactions within cells and communication between cell populations. Cell– cell communication (CCI) analysis can reveal potential signaling relationships among liver cell populations, while protein–protein interaction (PPI) networks provide a complementary molecular view of the interaction environments in which proteins function. Integrating these layers with disease- and cell-type-specific transcriptomic information could therefore provide a more comprehensive framework for investigating how cellular and molecular networks are remodeled across liver diseases. However, existing liver-focused resources still offer limited systematic exploration across cellular states, gene expression, intercellular communication, and protein interaction networks.

Here, we present **LiverHomo**, an integrated human liver single-cell resource constructed from 16 publicly available scRNA-seq datasets spanning healthy liver and major liver disease contexts, including MASH, ALD, cirrhosis, PBC, PSC, HCC, and iCCA. Following standardized quality control, harmonization, and cell-type annotation, LiverHomo comprises approximately 1.16 million high-quality cells and provides a unified framework for exploring cellular composition, gene expression, differential expression, and disease-associated transcriptional programs across cell types and disease states (Fig. 1A). Beyond transcriptomic analysis, LiverHomo incorporates CCI analysis and predicted context-specific PPI networks constructed using interaction information from our recently developed PrePPI-AF and ZEPPI frameworks (16,17), enabling exploration from cellular states and transcriptional alterations to intercellular communication and molecular interaction networks. These data are accessible through an interactive web portal integrating Atlas and Network Explorers, downloadable underlying data, and Ask LiverHomo, a natural-language interface designed to facilitate access to relevant analyses and visualizations (Fig. 1B). Together, LiverHomo provides a multi-scale resource for investigating cellular and molecular alterations across human liver health and disease.

**Figure 1.**
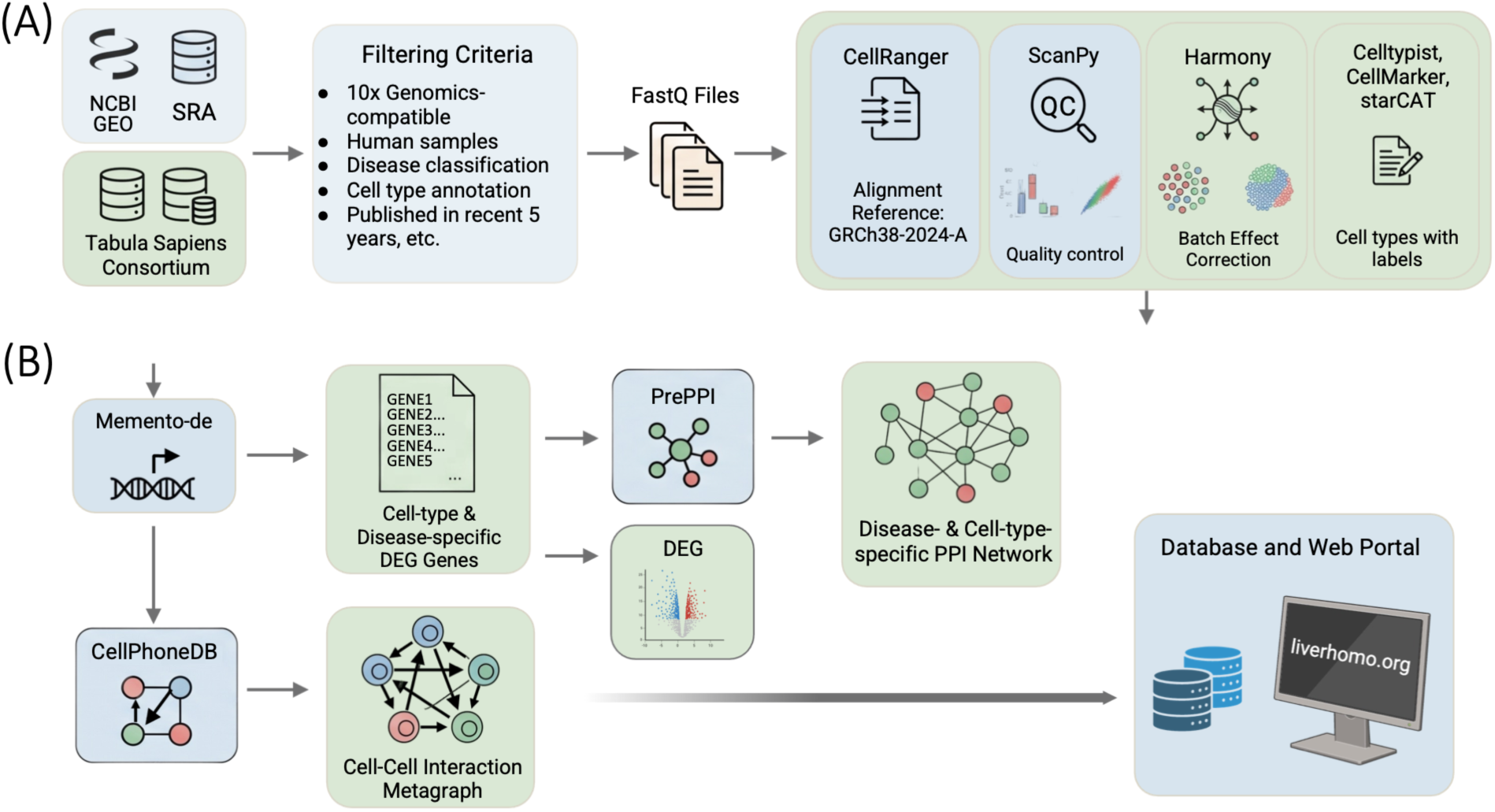
Data curation and preprocessing pipeline of LiverHomo. (A) We collected publicly available human liver scRNA-seq datasets from NCBI GEO, SRA, and the Tabula Sapiens consortium and curated them according to predefined criteria. Raw sequencing data were processed using Cell Ranger, followed by quality control with Scanpy, batch correction with Harmony, and cell-type annotation using CellTypist, CellMarker, and starCAT. (B) The harmonized single-cell atlas was used to characterize disease- and cell-type-specific molecular changes. Differential expression analysis with Memento identified context-specific upregulated genes, which were integrated with PrePPI to construct disease- and cell-type-specific PPI networks. In parallel, we used CellPhoneDB to infer disease-specific cell–cell interactions and construct intercellular communication networks.

## 2. Methods

### 2.1 Data collection, selection and curation

Human liver scRNA-seq datasets were identified through a systematic PubMed literature search, prioritizing studies on human liver tissues across healthy and diseased states. Datasets integrated into LiverHomo were sourced from: (i) the NCBI Gene Expression Omnibus (GEO) (20); (ii) the Sequence Read Archive (SRA) (https://www.ncbi.nlm.nih.gov/sra/docs/), guided by PubMed results; and (iii) the Tabula Sapiens consortium (21).

Publications were manually reviewed to curate sample metadata, including sequencing platform (restricted to 10x Genomics-compatible data), species (human only), disease classification, cell-type annotations and marker genes, publication details (published between 2019 and 2024), cell isolation methods, and library preparation protocols. Datasets with incomplete metadata or non-human samples were excluded to ensure atlas quality and consistency.

Raw FASTQ files were downloaded using the iSeq tool. Gene expression count matrices were generated with CellRanger (v7.1.0; 10x Genomics), aligned to the human reference genome refdata-gex-GRCh38-2024-A (Ensembl release 110). This standardized pipeline facilitated uniform preprocessing across all datasets.

### 2.2 Data processing

Gene expression matrices were analyzed using ScanPy (22) from OmicVerse with CPU GPU mixed acceleration (23) with standard quality control procedures applied to remove low-quality cells. Potential doublets were removed with Scrublet (24). Data were normalized and log-transformed. Batch effects were corrected using Harmony (25), followed by dimensionality-reduced embeddings using principal component analysis (PCA) and uniform manifold approximation and projection (UMAP) (26). Cell clustering was conducted using the Leiden algorithm with resolutions set at 0.1, 0.2, 0.3, 0.5, 0.7, 0.9, 1.0, and 1.2 (27).

### 2.3 Cell Type Annotation and Differential gene expression analysis

Cell type annotation was performed with both automated label transfer and manual curation. Initial annotations were obtained using CellTypist (28, 29), followed by validation through cluster-specific differentially expressed genes (DEGs) analysis and cross-referenced comparison with canonical lineage markers compiled from literature and the CellMarker database (30–36). Major liver cell populations were then annotated based on consensus between computational predictions and marker expression. T cells were further resolved using starCAT to capture subtype heterogeneity (37).

### 2.4 Inference of Cell-Cell Communication Networks

Cell-cell communication or interaction (CCI) was characterized by ligand-receptor interactions inferred by CellPhoneDB (38). Interactions were predicted in a cell type and disease specific context, retaining only genes expressed in at least a fraction of cells. Interaction scores were estimated from expression levels of cognate ligand-receptor pairs. The cell-type specificity of each interaction was assessed with permutation testing, and repeated subsampling was used to assess robustness. Consensus CCI networks were then constructed for each disease condition and cell type.

### 2.5 Cell-specific PPI Construction

We constructed cell-specific protein–protein interaction (PPI) networks for LiverHomo using the PPI resources and context-specific network construction strategy developed as part of the LiverDCP framework (39). Disease-associated differentially expressed genes were identified for each disease condition relative to healthy controls using Memento (40), while cell-type-specific marker genes were identified by comparing each cell type with all other cell populations in the healthy cohort. Protein interaction information was obtained from genome-wide PPI predictions of PrePPI-AF (16) and contextualized using the gene sets derived from the LiverHomo single-cell atlas. For each cell type, two complementary PPI subnetworks were constructed. A baseline cell-type-specific network was generated from interactions involving cell-type marker genes identified in healthy liver, representing the characteristic interaction environment of each cell population. Disease-specific networks were generated using genes upregulated in each disease condition to capture candidate disease-associated changes in the protein interaction environment (downregulated genes in the diseases are therefore not specifically represented). Interactions were further restricted to proteins with detectable expression in the corresponding cell type and prioritized according to PrePPI-AF likelihood-ratio scores and structural compatibility. The resulting disease- and cell-type-specific networks constitute the PPI layer presented through the LiverHomo Network Explorer. These networks represent expression-supported candidate subnetworks derived from predicted interactinos and do not indicate measured changes in physical binding or interaction strength.

### 2.6 Database and web portal implementation

#### Web architecture

The LiverHomo portal is implemented as a Python 3.11 web application using the Flask framework and deployed on a Linux server through the Gunicorn WSGI server. The backend uses AnnData, pandas, and NumPy for data retrieval and processing, Matplotlib and seaborn (18) for visualization, and NetworkX (19) for network layout computation. Web pages are rendered using server-side Jinja2 templates, while client-side JavaScript communicates with JSON-based REST endpoints to support interactive data retrieval and visualization without requiring full-page reloads.

The portal comprises four principal sections: a landing page summarizing the integrated atlas; the Atlas Explorer, which provides cell atlas visualization, gene expression analysis, and differential expression results (40); the Network Explorer, which presents context-specific PPI networks, Gene Ontology (GO) networks (41, 42), and cell–cell communication analyses; and the Resources page, which provides source dataset information and downloadable metadata. The Ask LiverHomo assistant is integrated into the landing and analysis pages to support natural-language data exploration.

#### Data organization and retrieval

The integrated LiverHomo atlas is represented as an AnnData (.h5ad) object containing approximately 1.16 million cells and 31,351 genes. To accommodate the memory and storage constraints of web hosting, we transformed the complete processed object, approximately 18 GB in size, into smaller, query-oriented data products for deployment. These include: (i) a metadata-only object containing per-cell annotations, disease classifications, source dataset information, and quality-control metrics; (ii) a condensed expression object containing log-normalized expression values and raw counts in compressed sparse row format; (iii) pre-rendered UMAP visualizations for supported metadata attributes; and (iv) a filter catalog defining the available disease and cell-type selections. Expression values are retrieved from HDF5 storage at the individual-gene level, avoiding the need to load the complete expression matrix into memory. Retrieved expression vectors are cached on disk to accelerate subsequent queries, while frequently accessed metadata are retained in memory as needed.

Precomputed analysis results are organized by disease condition and cell type. These include DEG tables, PrePPI-AF-derived interaction edge lists, CellPhoneDB interaction summaries, volcano plots, GO biological-process networks, and cell–cell communication heatmaps. A centralized configuration system manages file paths through to support consistent deployment across development and production environments. Gene queries are case-insensitive, with approximate symbol matching used to suggest alternatives when an exact match is unavailable.

#### Interactive visualization

LiverHomo provides six complementary visualization modules. (1) *Cell atlas visualization:* UMAP embeddings of the integrated atlas are displayed using pre-rendered images colored by supported metadata attributes, enabling rapid visualization of cellular distributions and annotations. (2) *Gene expression analysis:* Users can query individual genes and examine expression distributions through histograms and violin plots across selected disease conditions and cell populations. Figures are generated on demand, cached for repeated access, and downloadable in PNG format. (3) *Differential expression analysis:* Disease- and cell-type-specific DEG tables can be filtered by direction of expression change, sorted by individual attributes, and downloaded as CSV files. Corresponding volcano plots provide an overview of disease-associated transcriptional changes. (4) *Protein interaction networks:* Users can explore disease- and cell-type-specific PPI networks and restrict the displayed interactions to selected genes. Network layouts are generated using spring, Kamada–Kawai, or random positioning algorithms. Interactive SVG visualizations support zooming, panning, and figure export, while the underlying interaction edge lists are available for download. (5) *Gene Ontology networks:* Precomputed GO biological-process (41, 42) enrichment networks are presented for individual disease and cell-type contexts, enabling functional interpretation of associated gene sets. (6) *Cell–cell communication:* Disease-specific heatmaps summarize predicted interaction counts between cell populations. For selected cell-type pairs, comparative bar plots display interaction patterns across disease conditions. These modules provide complementary access to cellular composition, transcriptional variation, intercellular communication, and protein interaction networks within a unified web interface.

### 2.7. AI-assisted data exploration

LiverHomo incorporates *Ask LiverHomo*, a natural-language interface designed to facilitate access to the atlas and its associated analyses. The assistant uses the OpenAI GPT-5.4 nano model (gpt-5.4-nano) through the Chat Completions API with function calling. Rather than solely relying on model-generated biological interpretations, the assistant translates user questions into predefined server-side operations that retrieve data or generate visualizations directly from LiverHomo.

Five data-retrieval functions are implemented: (i) retrieving differential expression statistics for a queried gene across disease and cell-type contexts, including effect sizes and statistical significance; (ii) returning ranked DEG lists for selected diseases, cell types, and expression directions; (iii) summarizing gene expression across disease–cell-type groups using the proportion of expressing cells and mean log-normalized expression, restricted to groups containing at least 20 cells; (iv) retrieving cell counts by disease and cell type; and (v) providing information about the atlas, supported disease conditions, cell populations, and available analyses. An additional visualization function invokes the portal’s predefined plotting routines to generate gene expression distributions, volcano plots, UMAP visualizations, CCI heatmaps, and PPI networks.

To improve query interpretation, disease and cell-type terminology is standardized using a curated synonym dictionary, including mappings such as NASH to MASH and Kupffer cells to the broader macrophage population annotated in the atlas. Broader terms, such as T cells, can be mapped to the corresponding annotated subsets. Gene queries require an exact case-insensitive symbol match, with alternative symbols suggested when no match is found.

The assistant supports multi-step queries through a maximum of five rounds of function calling and retains the six most recent conversation messages to enable contextual follow-up questions. A system-level instruction constrains responses to the atlas and its supported analyses. When relevant general biological background is needed to answer user’s question, the answer will begin with “Background (general knowledge):” to distinguish from atlas-derived information. Requests outside the scope of LiverHomo, including medical diagnosis or treatment advice, will receive a predefined response indicating that the question falls outside the resource’s scope.

## 3. Results

### 3.1 Overview of the LiverHomo Resource

LiverHomo integrates scRNA-seq data from more than 350 individuals across 16 publicly available datasets, comprising 1,157,134 high-quality cells spanning healthy and diverse liver disease contexts. The integrated atlas provides a foundation for exploring cellular heterogeneity, disease-associated transcriptional changes, cell–cell communication, and context-specific protein interaction networks.

The LiverHomo web portal is organized into four main sections: Home, Atlas Explorer, Network Explorer, and Resources. The Home page summarizes the resource, including dataset composition, cell counts, disease coverage, and access to the principal analytical modules. The Atlas Explorer lets users investigate cellular distributions, gene expression patterns, and differential gene expression across diseases and cell types. The Network Explorer extends these analyses to molecular and intercellular relationships through cell–cell communication networks, context-specific protein–protein interaction (PPI) networks, and Gene Ontology (GO)-annotated (41, 42) interaction networks. The Resources page provides information on source datasets, data processing, analytical methods, and data access. Users can also download data tables and visualization outputs directly from the corresponding analytical modules, supporting further investigation and reuse.

### 3.2 Interactive Data Exploration and Visualization

#### Atlas Explorer

The Atlas Explorer provides interactive access to the cellular and transcriptional landscapes represented in the integrated LiverHomo atlas. UMAP visualizations (26) allow users to examine cellular distributions across disease conditions and cell types. Users can select one or multiple disease states and cell populations to explore their distributions within the atlas and retrieve associated cell-level metadata, including cell complexity, disease classification, source dataset, and cell-type annotations. Selected visualizations and metadata are available for download.

The Gene Expression module enables users to investigate the expression of individual genes across selected disease conditions and cell populations (Fig. 2A). Users can select multiple diseases and cell types simultaneously, allowing comparative visualization through overlapping histograms or grouped violin plots. Histograms show expression distributions for selected cell populations across disease conditions, whereas violin plots organize expression profiles by disease and cell type, enabling comparison of expression heterogeneity across biological contexts.

**Figure 2.**
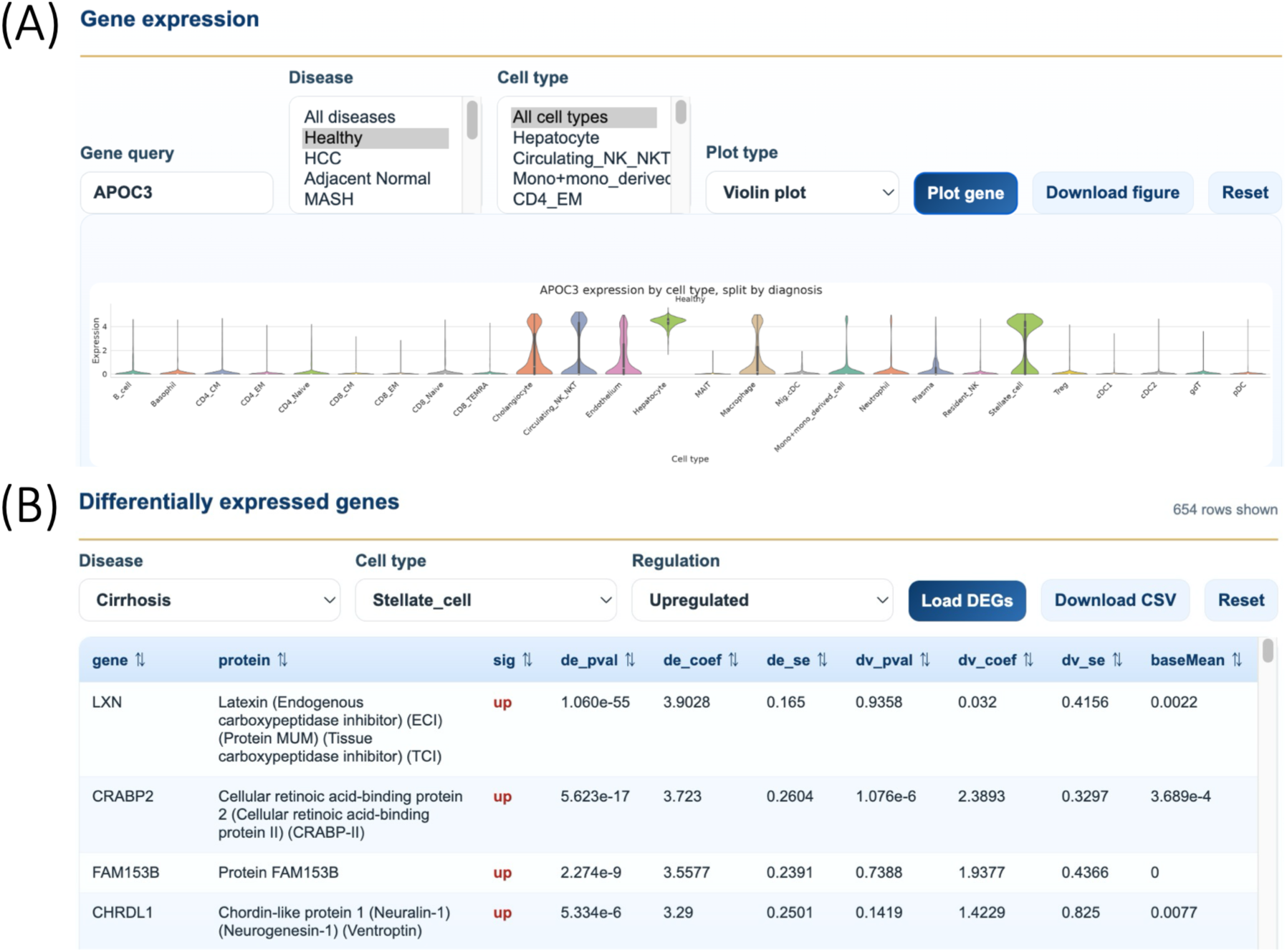
Interactive exploration of the LiverHomo single-cell atlas. (A) Gene Expression module demonstrating the visualization of APOC3 expression across cell types in healthy liver using violin plots. (B) Differential Expression module displaying upregulated genes in hepatic stellate cells from cirrhotic liver relative to healthy controls.

The Differential Expression module provides access to disease-associated transcriptional changes through searchable DEG tables and volcano plots. Users can select disease and cell-type contexts, filter genes by the direction of expression change, and sort results according to statistical significance or effect size (Fig. 2B). The tables include gene identifiers, encoded proteins, differential expression coefficients, standard errors, and measures of statistical significance. Gene-to-protein annotations were obtained from reviewed UniProt records. The underlying DEG tables and corresponding visualizations such as volcano plots are available for download.

#### Network Explorer

The Network Explorer extends LiverHomo beyond cellular and transcriptional states by providing access to context-specific protein interactions, functional pathways, and intercellular communication networks (Fig. 3). The PPI module enables users to explore predicted protein interaction networks across liver diseases and cell types. Users can select a biological context and further focus the network on one or more genes of interest. For selected genes, the viewer extracts their one-hop interaction neighborhoods, allowing examination of their immediate protein interaction environments (Fig. 3A). Interactive network visualizations support alternative layouts, node-degree-based representations, zooming, and panning. Users can export both the displayed networks and underlying interaction tables for further analysis.

**Figure 3.**
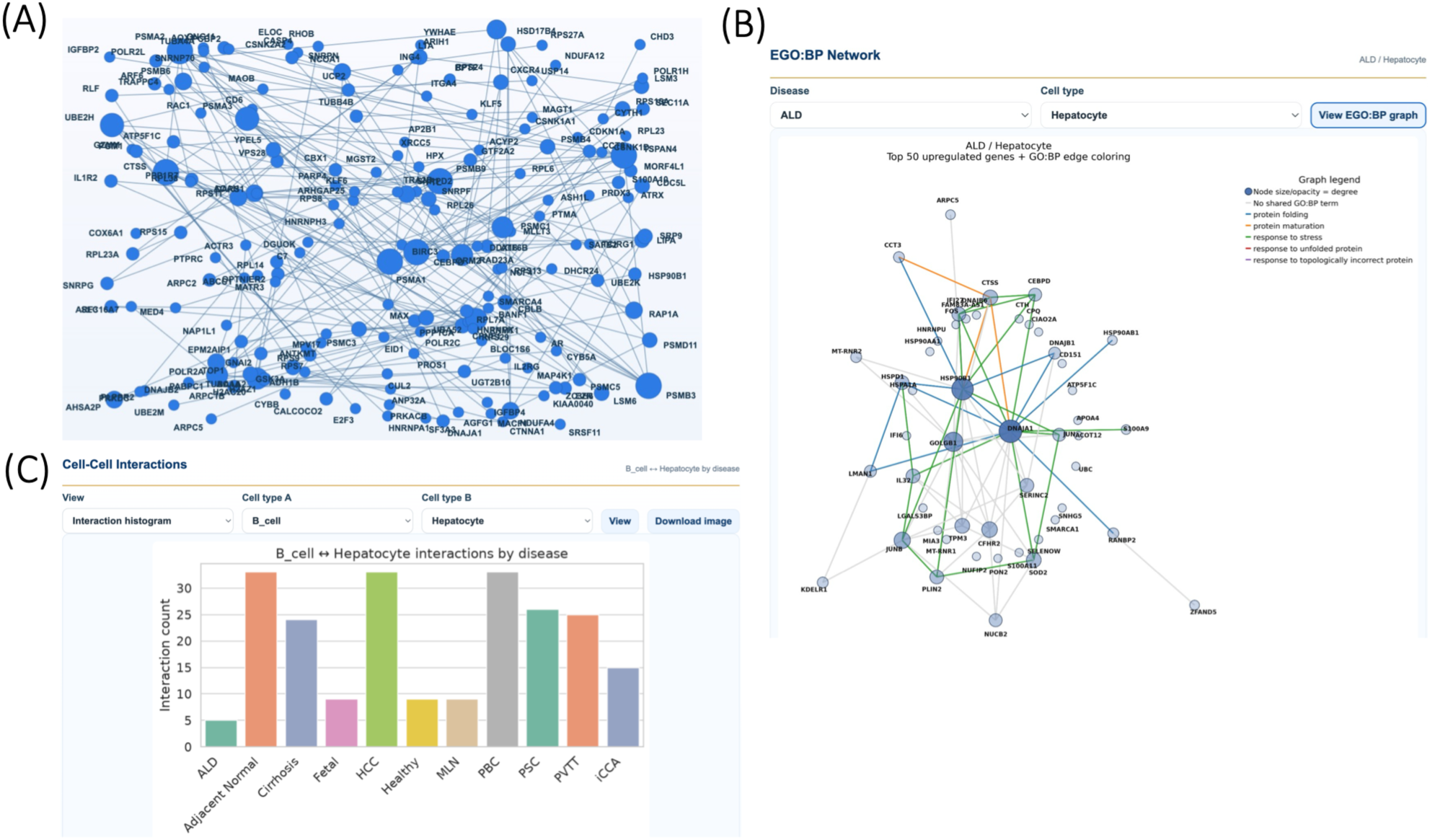
Interactive exploration of the LiverHomo Network Explorer. (A) Context-specific protein–protein interaction (PPI) network for regulatory T cells (Tregs) in metabolic dysfunction-associated steatohepatitis (MASH), displaying 193 nodes and the top 200 interactions ranked by PrePPI score from 149,389 predicted interactions, using a random network layout. (B) Gene Ontology biological process (GO:BP)-annotated PPI network for hepatocytes in alcohol-associated liver disease (ALD). (C) Comparison of predicted cell–cell interactions between B cells and hepatocytes across liver disease conditions.

To facilitate functional interpretation, the GO-annotated PPI module displays interaction networks associated with enriched biological processes (Fig. 3B). For each disease–cell-type context, we construct networks from the top 50 upregulated genes ranked by differential expression coefficient and annotated using the five highest-ranked GO biological-process terms. For disease-level networks combining all cell types, the analysis includes up to 200 genes and 20 GO terms. These visualizations highlight interactions among proteins associated with shared biological processes and provide a functional perspective on disease-associated PPI networks.

The Cell–Cell Communication module provides disease-specific representations of predicted ligand–receptor interactions between liver cell populations. Heatmaps summarize interaction counts between cell types using a consistent color scale across diseases (Fig. 3C). Because these counts depend on cell populations and cell counts sampled in each condition, cross disease differences should be interpreted decriptively. Chord diagrams provide complementary visualizations of communication patterns within individual disease contexts. Users can also select a pair of cell types to compare their predicted communication across diseases through bar plots. (Fig, 3C) These visualizations are available for download.

#### Ask LiverHomo

Beyond conventional navigation through its analytical modules, LiverHomo provides Ask LiverHomo, a natural-language interface for querying atlas data and generating visualizations. Users can explore gene expression, differential expression, cell-type distributions, and network relationships without manually navigating or configuring individual analysis modules (Fig. 4). The assistant interprets user queries and invokes predefined server-side functions to retrieve atlas-derived results or generate corresponding visualizations, using the same underlying data retrieval and analytical procedures as the portal. By allowing users to directly describe their biological questions, Ask LiverHomo reduces the need to familiarize themselves with the portal’s structure and available analytical functions, lowering the learning curve and facilitating more intuitive and efficient data exploration. The assistant also supports multilingual and mixed-language queries, further improving accessibility for users with diverse linguistic backgrounds and work habits.

**Figure 4.**
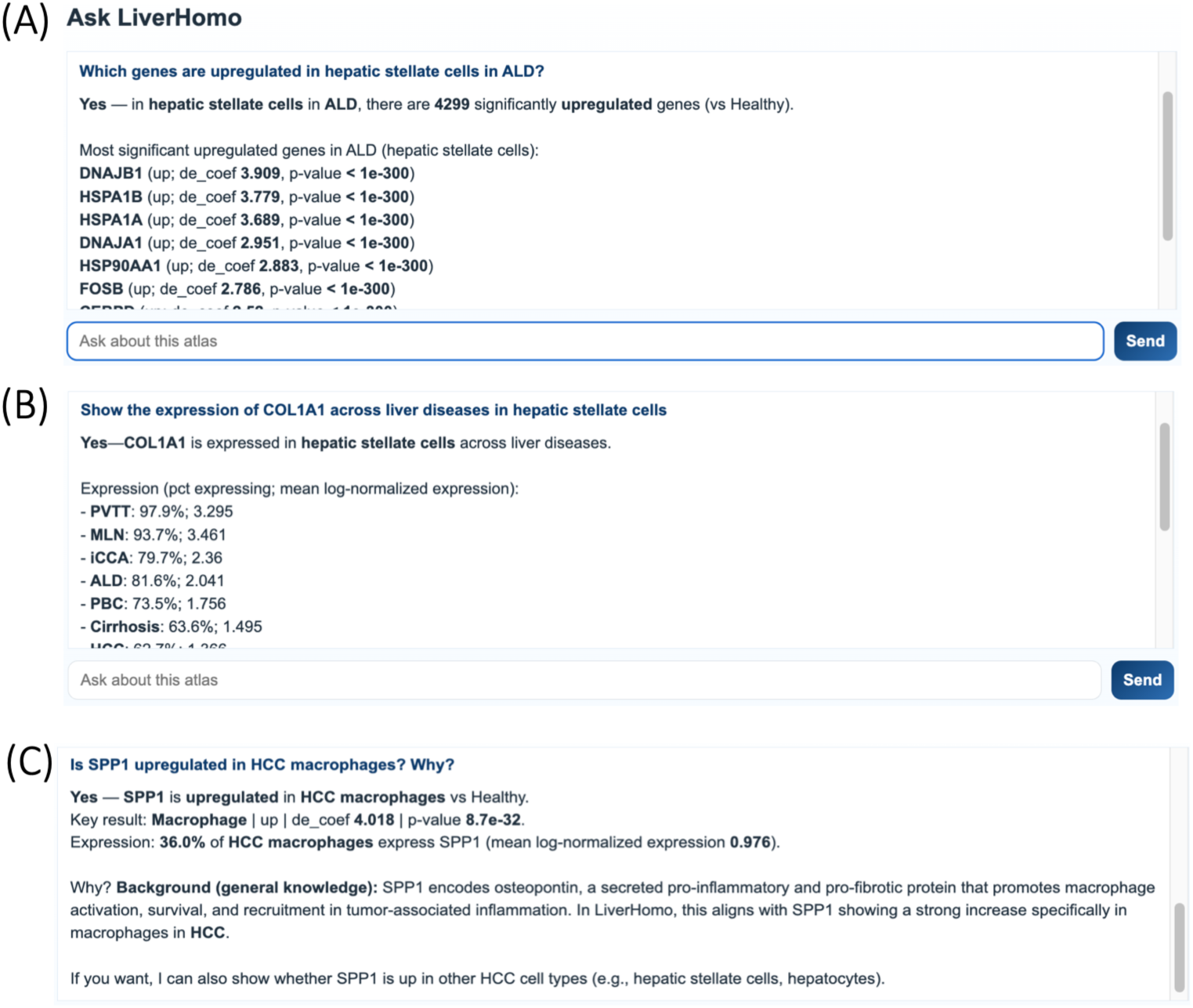
Interactive exploration of LiverHomo using the Ask LiverHomo AI assistant. (A) Identification of differentially expressed genes in hepatic stellate cells from ALD. (B) Natural-language query requesting visualization of COL1A1 expression across liver disease conditions in hepatic stellate cells. (C) Query examining whether SPP1 is upregulated in macrophages from HCC, with the assistant providing atlas-derived differential expression results and relevant biological background.

### 3.3 Biological Use Cases

#### Context-dependent NFE2L2 expression and interaction networks in MASH

MASH (metabolic dysfunction-associated steatohepatitis) is characterized by hepatic steatosis, inflammation, and hepatocellular injury, with the potential to progress to fibrosis and cirrhosis (43). Oxidative stress is a major contributor to MASH pathogenesis, promoting lipid peroxidation, mitochondrial dysfunction, inflammatory responses, and hepatocellular injury (44–48). NFE2L2 encodes the transcription factor NRF2 and plays a central role in cellular antioxidant defense, redox homeostasis, and protection against oxidative stress-induced damage (49–54).

To illustrate LiverHomo’s utility for investigating disease-associated molecular changes, we examined NFE2L2 expression and its predicted protein interaction network across liver conditions. GO-annotated PPI analysis of naive CD8+ T cells placed NFE2L2 in a network associated with biological processes including apoptotic signaling and cellular responses to stress (Fig. 5A). Differential expression analysis showed increased NFE2L2 expression in MASH compared with healthy liver (Fig. 5B), consistent with a potential transcriptional response to disease-associated cellular stress, although transcript abundance alone does not establish NRF2 protein stabilization or activity. Interestingly, NFE2L2 expression was lower in MASH than in adjacent normal liver tissue (Fig. 5C), indicating that the direction of observed difference depends on the reference tissue context. These observations illustrate how LiverHomo can integrate differential expression and functional network annotations to investigate context-dependent molecular changes associated with liver disease.

**Figure 5.**
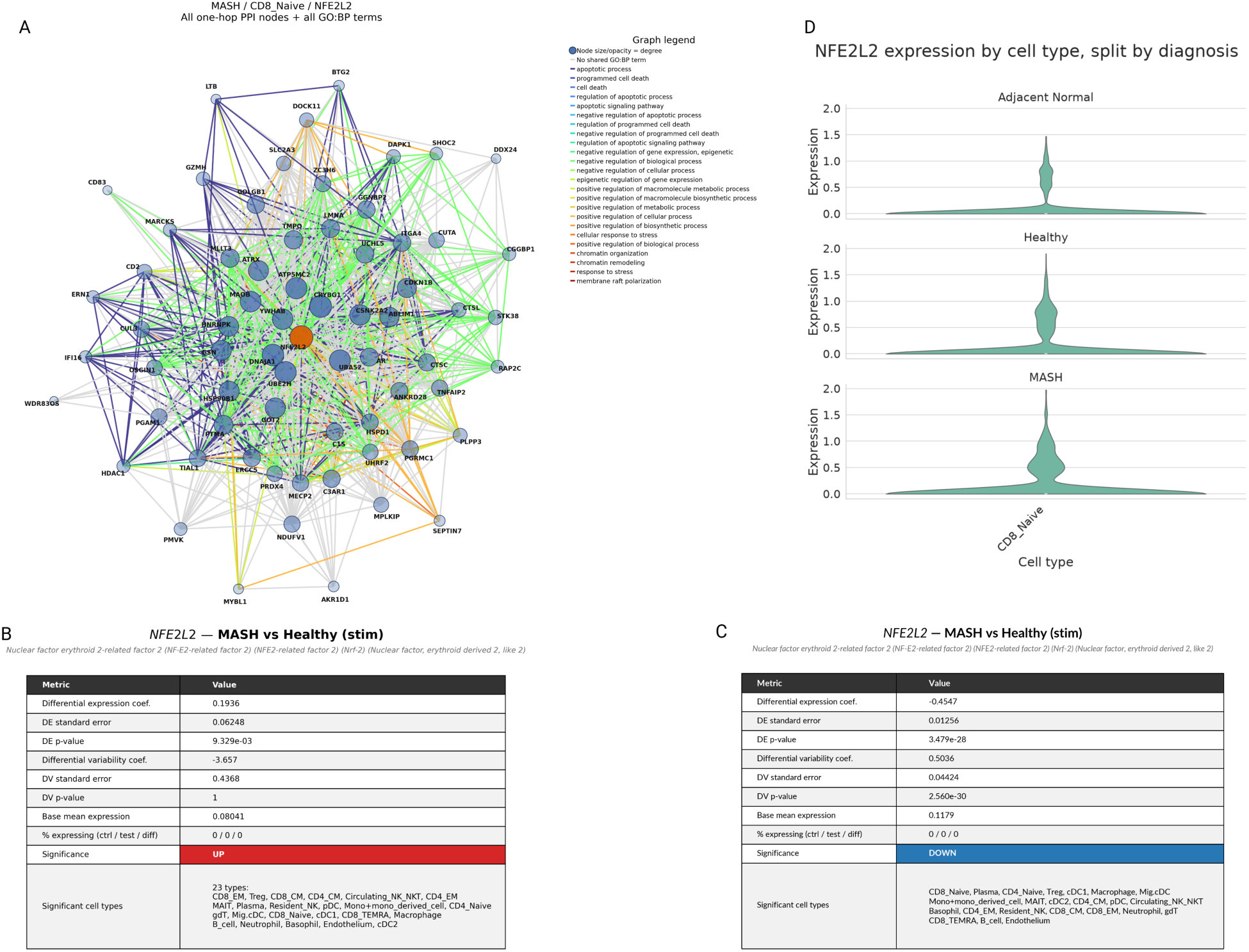
Context-dependent NFE2L2 expression and predicted protein interaction networks in MASH. (A) Gene Ontology biological process (GO:BP)-annotated PPI network for naive CD8+ T cells in MASH highlighting NFE2L2 and its association with biological processes related to apoptotic signaling and cellular stress responses. (B) Differential expression analysis showing increased NFE2L2 expression in metabolic dysfunction-associated steatohepatitis (MASH) compared with healthy liver. (C) Differential expression analysis showing decreased NFE2L2 expression in MASH compared with adjacent-normal liver tissue. (D) Violin plots showing the relative gene expression level of NFE2L2 in health, adjacent normal, and MASH cells, matching the previous tables.

#### Altered Expression of the Circadian Regulator CLOCK in HCC

Circadian rhythm regulation has emerged as an important factor in cancer development and progression. CLOCK and BMAL1 (encoded by ARNTL) form a transcriptional regulatory complex that coordinates circadian gene expression and influences cellular processes involved in metabolism, cell-cycle regulation, and DNA damage responses (55, 56). Multiple mice studies have shown that mutant circadian genes or chronic circadian rhythm disruptions lead to increased risk for HCC (57–59). Multiple mechanisms have been shown linking these two conditions together. Chronic jet leg disrupts sympathetic function, driving constitutive androstane receptor (CAR) and MAPK-dependent AP-1 activation. AP-1 then abnormally upregulates c-Myc, promoting cell growth and survival, thus leading to tumorigenesis (57, 60). Circadian disruption can also blunt PER1-mediated activation of ATM and alter CLOCK-BMAL1-dependent SIRT1-mediated acetylation of p53. Loss of ATM and p53 activity results in defective apoptosis, impaired cell cycle arrest, and reduced DNA repair, leading to increased tumor growth and genomic instability (60, 61). More specifically, the downregulation of CLOCK/BMAL1 has been shown to arrest HCC in G2/M phase of the cell cycle while its upregulation drives the proliferation of HCC (62).

Using the LiverHomo Differential Expression module, we examined CLOCK expression in hepatocytes across healthy and HCC liver tissues. CLOCK was significantly upregulated in hepatocytes from HCC tissues compared with healthy controls (Fig. 6A), suggesting that circadian rhythm plays a role in HCC pathogenesis. This is consistent with previously reported associations between circadian regulators and HCC biology. UPon further investigation of the data, MYC which encodes for c-Myc and SIRT1 were also found to be significantly upregulated in HCC cells (Fig. 6B and Fig.6C), matching the previously described literature. This example demonstrates how LiverHomo enables cell-type-resolved investigation of disease-associated transcriptional changes and facilitates the exploration of biologically relevant genes across liver disease contexts.

**Figure 6.**
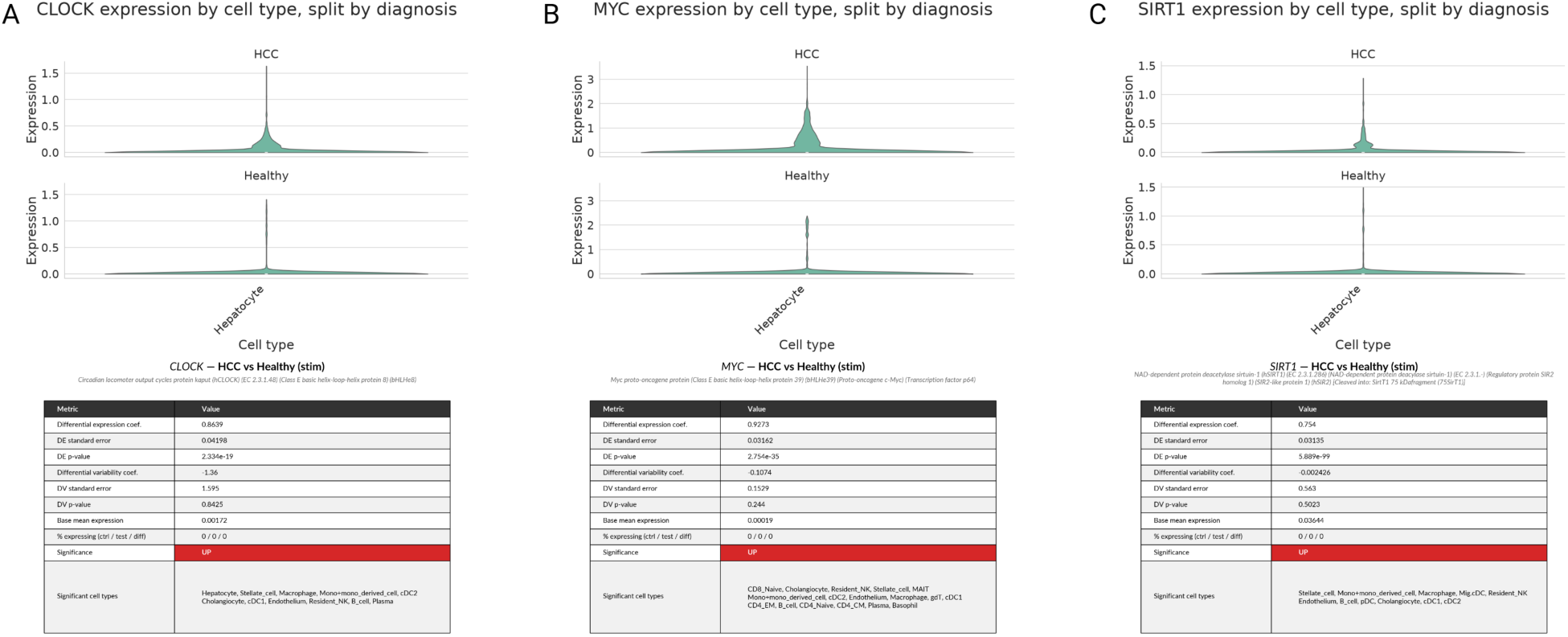
Context-dependent expression of Significant Circadian Regulated Genes in HCC. (A) Differential expression analysis showing increased CLOCK expression in hepatocellular carcinoma (HCC compared with healthy liver (B) Differential expression analysis showing increased MYC expression in hepatocellular carcinoma (HCC compared with healthy liver. (C) Differential expression analysis showing increased SIRT1 expression in hepatocellular carcinoma (HCC) compared with healthy liver.

### 3.4 Comparisons with Existing Liver Resources

To contextualize LiverHomo among existing liver-focused resources, we compared its data coverage and analytical capabilities with six publicly available databases: LiverSCA (63), GepLiver (64), CancerLivER (65), Liver Cell Atlas (66), Liver Single Cell Atlas (67), and scAtlasLC (68). These resources provide complementary functionalities for investigating liver biology, including gene expression visualization, differential expression analysis, cell–cell communication, functional enrichment, and cancer-associated transcriptional changes. Several databases offer specialized analytical capabilities, such as survival analysis in GepLiver, trajectory analysis in LiverSCA, and comparisons of gene expression between malignant and nonmalignant cells in scAtlasLC.

LiverHomo complements these resources by integrating harmonized single-cell transcriptomic data with cell–cell communication and context-specific PPI networks across healthy and multiple liver disease conditions. In addition to conventional gene expression and differential expression analyses, LiverHomo provides access to predicted PPI networks contextualized by disease and cell type, together with Gene Ontology annotations for functional interpretation. These integrated data layers enable users to investigate disease-associated transcriptional changes alongside predicted molecular and intercellular interactions within a unified resource.

Another feature of LiverHomo is Ask LiverHomo, a natural-language interface that enables users to retrieve atlas-derived results and generate visualizations through predefined analytical functions. This interface offers an alternative to conventional module-based navigation and facilitates intuitive access to the resource without requiring programming expertise. Together, the integration of transcriptomic and interaction-network data, broad liver disease representation, and multiple data-access interfaces distinguish LiverHomo as a complementary resource for exploring cellular and molecular changes in human liver health and disease.

## 4. Discussion

### 4.1 Applications and Limitations

LiverHomo provides an integrated resource for investigating cellular and molecular changes across healthy and diseased human liver tissues. By harmonizing publicly available single-cell transcriptomic datasets and incorporating complementary network-level analyses, LiverHomo enables researchers to explore disease-associated gene expression, cellular heterogeneity, cell–cell communication, and context-specific protein interactions within a unified framework. The interactive web portal and Ask LiverHomo assistant further facilitate access to these data without requiring extensive computational expertise.

The Gene Expression and Differential Expression modules enable systematic comparisons of transcriptional profiles across liver diseases and cell populations, supporting the identification of candidate disease-associated genes and potential biomarkers for further investigation. The Cell– Cell Communication module provides complementary insights into predicted ligand–receptor interactions between cell populations, helping explore intercellular signaling changes associated with disease progression. The Network Explorer extends these analyses to predicted protein interaction networks, allowing users to examine the molecular interaction environments of genes of interest and their associations with biological processes. Together, these complementary analyses support hypothesis generation and prioritize molecular and cellular mechanisms for subsequent experimental validation.

Several limitations should be considered when using LiverHomo. First, individual cell populations vary across disease conditions, reflecting differences in the experimental designs, tissue sampling, and cell-isolation procedures of the original studies. This is particularly evident in MASH, where hepatocytes are largely absent from the available scRNA-seq datasets. For example, GSE169446 specifically enriched dendritic cells, whereas GSE159977 focused on hepatic immune-cell populations, particularly T cells. These experimental designs preferentially captured non-parenchymal cells, limiting hepatocyte representation despite their abundance in liver tissue. In addition, the large size and fragility of hepatocytes can complicate their recovery using conventional scRNA-seq protocols. Thus, the observed cell-type distributions reflect the sampling strategies of the source studies and should not be interpreted as direct estimates of cellular abundance in liver tissue. Future incorporation of additional whole-cell or single-nucleus RNA-seq datasets may improve hepatocyte coverage and enable more comprehensive analyses of MASH-associated transcriptional changes.

Second, transcriptomic measurements do not necessarily reflect protein abundance or activity. Accordingly, the context-specific PPI networks, constructed from computationally predicted protein interactions and cell-type- and disease-associated transcriptomic profiles, should be interpreted as representations of potential molecular relationships rather than experimentally validated interaction networks. Similarly, cell–cell communication networks inferred from ligand– receptor expression represents predicted signaling relationships that require experimental validation. These considerations are important when using LiverHomo to generate biological hypotheses or prioritize candidates for downstream investigation.

### 4.2 Future Directions

Future development of LiverHomo will focus on expanding disease and cell-type coverage and incorporating additional molecular data modalities. One priority is the integration of single-nucleus RNA sequencing (snRNA-seq) datasets, which may better represent hepatocytes and other cell populations underrepresented in conventional scRNA-seq studies. This expansion will be particularly valuable for MASH, where the current atlas has limited hepatocyte and hepatic stellate cell coverage, as well as for other disease conditions with incomplete cellular representation. Integrating these datasets will require careful harmonization of single-cell and single-nucleus transcriptomic profiles to account for differences in transcript detection and cell-type representation.

Another priority is to strengthen the connection between transcriptomic changes and protein-level molecular interactions. The current framework uses gene expression profiles to contextualize computationally predicted PPI networks; however, transcript abundance does not necessarily reflect protein abundance or interaction activity. Incorporating quantitative proteomics and spatially resolved protein measurements, including immunofluorescence-based approaches, could provide complementary evidence for evaluating disease-associated protein expression and refining context-specific interaction networks. Together, these extensions would improve the biological resolution of LiverHomo and support more comprehensive investigations of the molecular and cellular mechanisms underlying liver disease.

Beyond data exploration, LiverHomo may also serve as a foundation for developing computational models that integrate molecular interactions with cellular and disease contexts. For example, our recently developed LiverDCP framework (39) leverages liver single-cell transcriptomic profiles and context-specific interaction networks to learn disease- and cell-type-aware protein representations, supporting downstream applications such as disease-associated gene prioritization and therapeutic target identification. Future development could strengthen the connection between LiverHomo and LiverDCP, enabling users to move from interactive data exploration toward computational prediction and hypothesis generation.

## 5. Conclusions

LiverHomo provides an integrated resource for exploring cellular and molecular changes across healthy and diseased human liver tissues. By combining a harmonized single-cell transcriptomic atlas with differential expression analysis, cell–cell communication predictions, and context-specific protein–protein interaction networks, LiverHomo enables investigation of liver disease biology across multiple levels of organization. Its interactive visualization modules and Ask LiverHomo AI assistant facilitate data exploration and accessibility, supporting the identification of candidate biomarkers, prioritization of potential therapeutic targets, and generation of testable hypotheses. LiverHomo provides a foundation for future investigations of the cellular and molecular mechanisms underlying human liver health and disease.

## Data availability

The single-cell RNA-sequencing datasets integrated into LiverHomo were obtained from publicly available repositories, including the NCBI GEO, SRA, and Tabula Sapiens. Accession numbers, original publications, and associated metadata for all 16 source datasets are provided on the Resources page of the LiverHomo web portal (https://www.liverhomo.org/resources).

The harmonized LiverHomo atlas, comprising 1,157,134 high-quality cells from healthy and diseased human liver tissues, is accessible through the LiverHomo web portal (https://www.liverhomo.org). The portal provides interactive access to cell-type annotations, gene expression profiles, differential expression results, predicted cell–cell communication networks, and context-specific protein–protein interaction networks. Derived data tables and visualization outputs can be downloaded directly from the corresponding analytical modules. The curated cell-level metadata for all 1,157,134 cells, including cell-type annotations, disease conditions, source datasets, and quality-control metrics, are available for download in AnnData (.h5ad) format. Precomputed PPI and cell-cell communication datasets are publicly available through Zenodo (https://zenodo.org/records/20431227) as part of the LiverDCP training data.

## Code availability

The source code for the LiverHomo web portal, including its backend data retrieval functions, interactive visualization modules, and Ask LiverHomo AI assistant, is publicly available on GitHub at https://github.com/zhaolabutmb/LiverHomo. The repository includes documentation and instructions for installation and usage. Scripts used for data preprocessing, integration, cell-type annotation, differential expression analysis, and network construction are available in the same repository.

## Acknowledgements

We thank Dr. Erez Lieberman Aiden and Caleb Chiao for insightful discussions on AI tools, and Drs. Clem Wong and John Perkyns of the SCSB Scientific Computing team for their technical support in maintaining our computing infrastructure. This work was supported by the Welch Foundation (H-2308-20260402) and the UT System Rising STARs Award (to H.Z.).

## Author contributions

H.Z., S.Y., and Z.S. conceived and designed the research; S.Y., Z.S., Zh.S., and H.Z. performed the research; S.Y. and H.Z. designed the web portal; S.Y., Z.S., E.A.S., O.S., and H.S. analyzed the data; D.G., and B.D. contributed to the biological applications; S.Y., Z.S., and H.Z. wrote the manuscript. All authors reviewed and revised the manuscript.

## References

1. Llovet JM, Kelley RK, Villanueva A, Singal AG, Pikarsky E, Roayaie S, et al. Hepatocellular carcinoma. Nat Rev Dis Primer [Internet]. 2021 Jan 21 [cited 2025 Apr 24];7(1):1–28. Available from: https://www.nature.com/articles/s41572-020-00240-3

2. F L, W Z, F Z, Y X, X H, G J, et al. DAB2 + macrophages support FAP + fibroblasts in shaping tumor barrier and inducing poor clinical outcomes in liver cancer. Theranostics [Internet]. 2024 Aug 12 [cited 2024 Oct 14];14(12). Available from: https://pubmed.ncbi.nlm.nih.gov/39239526/

3. Ho DWH, Tsui YM, Chan LK, Sze KMF, Zhang X, Cheu JWS, et al. Single-cell RNA sequencing shows the immunosuppressive landscape and tumor heterogeneity of HBV-associated hepatocellular carcinoma. Nat Commun [Internet]. 2021 June 17 [cited 2025 Feb 12];12:3684. Available from: https://www.ncbi.nlm.nih.gov/pmc/articles/PMC8211687/

4. D’souza S, Lau KC, Coffin CS, Patel TR. Molecular mechanisms of viral hepatitis induced hepatocellular carcinoma. World J Gastroenterol. 2020 Oct 14;26(38):5759–83.

5. Rumgay H, Arnold M, Ferlay J, Lesi O, Cabasag CJ, Vignat J, et al. Global burden of primary liver cancer in 2020 and predictions to 2040. J Hepatol [Internet]. 2022 Dec [cited 2025 Sept 1];77(6):1598–606. Available from: https://www.ncbi.nlm.nih.gov/pmc/articles/PMC9670241/

6. Sakata M, Kitada K, Omote R, Sonobe H, Utsumi M, Tokunaga N, et al. Synchronous double primary combined hepatocellular-cholangiocarcinoma and cholangiolocarcinoma in a cirrhotic liver. J Clin Transl Hepatol [Internet]. 2023 Aug 28 [cited 2025 Sept 1];11(4):991–7. Available from: https://www.ncbi.nlm.nih.gov/pmc/articles/PMC10318272/

7. Dash S, Aydin Y, Widmer KE, Nayak L. Hepatocellular carcinoma mechanisms associated with chronic HCV infection and the impact of direct-acting antiviral treatment. J Hepatocell Carcinoma [Internet]. 2020 Apr 15 [cited 2025 Sept 1];7:45–76. Available from: https://www.ncbi.nlm.nih.gov/pmc/articles/PMC7167284/

8. Sirica AE, Strazzabosco M, Cadamuro M. Intrahepatic cholangiocarcinoma: Morpho-molecular pathology, tumor reactive microenvironment, and malignant progression. Adv Cancer Res [Internet]. 2021 [cited 2025 Sept 1];149:321–87. Available from: https://www.ncbi.nlm.nih.gov/pmc/articles/PMC8800451/

9. Younossi ZM, Kalligeros M, Henry L. Epidemiology of metabolic dysfunction-associated steatotic liver disease. Clin Mol Hepatol [Internet]. 2025 Feb [cited 2025 Sept 1];31(Suppl):S32–50. Available from: https://www.ncbi.nlm.nih.gov/pmc/articles/PMC11925440/

10. Ying F, Chan MSM, Lee TKW. Cancer-associated fibroblasts in hepatocellular carcinoma and cholangiocarcinoma. Cell Mol Gastroenterol Hepatol [Internet]. 2023 Jan 25 [cited 2025 Sept 1];15(4):985–99. Available from: https://www.ncbi.nlm.nih.gov/pmc/articles/PMC10040968/

11. Xue R, Chen L, Zhang C, Fujita M, Li R, Yan SM, et al. Genomic and transcriptomic profiling of combined hepatocellular and intrahepatic cholangiocarcinoma reveals distinct molecular subtypes. Cancer Cell [Internet]. 2019 June 10 [cited 2025 Sept 1];35(6):932–947.e8. Available from: https://www.sciencedirect.com/science/article/pii/S1535610819302028

12. Kang J, Lee JH, Cha H, An J, Kwon J, Lee S, et al. Systematic dissection of tumor-normal single-cell ecosystems across a thousand tumors of 30 cancer types. Nat Commun [Internet]. 2024 May 14 [cited 2024 July 4];15(1):4067. Available from: https://www.nature.com/articles/s41467-024-48310-4

13. Liu K, Cui Y, Han H, Guo E, Shi X, Xiong K, et al. Fibroblast atlas: Shared and specific cell types across tissues. Sci Adv. 2025 Apr 4;11(14):eado0173.

14. Pan Q, Li B, Lin D, Miao YR, Luo T, Yue T, et al. scLiverDB: A database of human and mouse liver transcriptome landscapes at single-cell resolution. Small Methods. 2023 Sept;7(9):e2201421.

15. Long R, Suoangbaji, Ng IOL, Ho DWH. LiverSCA: A comprehensive and user-friendly cell atlas in human hepatocellular carcinoma. Comput Struct Biotechnol J [Internet]. 2024 June 24 [cited 2025 Feb 19];23:2740–5. Available from: https://www.ncbi.nlm.nih.gov/pmc/articles/PMC11266871/

16. Zhao H, Petrey D, Murray D, Honig B. ZEPPI: Proteome-scale sequence-based evaluation of protein-protein interaction models. Proc Natl Acad Sci U S A. 2024 May 21;121(21):e2400260121.

17. Petrey D, Zhao H, Trudeau SJ, Murray D, Honig B. PrePPI: A structure informed proteome-wide database of protein-protein interactions. J Mol Biol. 2023 July 15;435(14):168052.

18. Waskom, M. L., (2021). seaborn: statistical data visualization. Journal of Open Source Software, 6(60), 3021, 10.21105/joss.03021.

19. Aric A. Hagberg, Daniel A. Schult and Pieter J. Swart, “Exploring network structure, dynamics, and function using NetworkX”, in Proceedings of the 7th Python in Science Conference (SciPy2008), Gäel Varoquaux, Travis Vaught, and Jarrod Millman (Eds), (Pasadena, CA USA), pp. 11–15, Aug 2008

20. Barrett T, Wilhite SE, Ledoux P, Evangelista C, Kim IF, Tomashevsky M, et al. NCBI GEO: Archive for functional genomics data sets—update. Nucleic Acids Res [Internet]. 2013 Jan 1 [cited 2025 Sept 1];41(D1):D991–5. Available from: 10.1093/nar/gks1193

21. Tabula Sapiens Consortium*, Jones RC, Karkanias J, Krasnow MA, Pisco AO, Quake SR, et al. The tabula sapiens: A multiple-organ, single-cell transcriptomic atlas of humans. Science. 2022 May 13;376(6594):eabl4896.

22. Wolf FA, Angerer P, Theis FJ. SCANPY: Large-scale single-cell gene expression data analysis. Genome Biol [Internet]. 2018 Feb 6 [cited 2025 Apr 24];19(1):15. Available from: 10.1186/s13059-017-1382-0

23. Zeng Z, Ma Y, Hu L, Tan B, Liu P, Wang Y, et al. OmicVerse: A framework for bridging and deepening insights across bulk and single-cell sequencing. Nat Commun [Internet]. 2024 July 16 [cited 2025 Sept 1];15(1):5983. Available from: https://www.nature.com/articles/s41467-024-50194-3

24. Wolock SL, Lopez R, Klein AM. Scrublet: Computational identification of cell doublets in single-cell transcriptomic data. Cell Syst [Internet]. 2019 Apr 24 [cited 2025 Sept 1];8(4):281–291.e9. Available from: https://www.ncbi.nlm.nih.gov/pmc/articles/PMC6625319/

25. Korsunsky I, Millard N, Fan J, Slowikowski K, Zhang F, Wei K, et al. Fast, sensitive and accurate integration of single-cell data with harmony. Nat Methods [Internet]. 2019 Dec [cited 2025 Apr 24];16(12):1289–96. Available from: https://www.nature.com/articles/s41592-019-0619-0

26. McInnes L, Healy J, Melville J. UMAP: Uniform manifold approximation and projection for dimension reduction [Internet]. arXiv; 2020 [cited 2025 Apr 24]. Available from: http://arxiv.org/abs/1802.03426

27. Traag VA, Waltman L, van Eck NJ. From louvain to leiden: Guaranteeing well-connected communities. Sci Rep [Internet]. 2019 Mar 26 [cited 2025 Apr 24];9(1):5233. Available from: https://www.nature.com/articles/s41598-019-41695-z

28. Automatic cell-type harmonization and integration across Human Cell Atlas datasets. Cell 186, 5876–5891.e20 (2023) Availabe from: https://www.cell.com/cell/fulltext/S0092-8674(23)01312-0?_returnURL=https%3A%2F%2Flinkinghub.elsevier.com%2Fretrieve%2Fpii%2FS0092867423013120%3Fshowall%3Dtrue

29. Cross-tissue immune cell analysis reveals tissue-specific features in humans. Science 376, eabl5197 (2022). Available from: https://www.science.org/doi/10.1126/science.abl5197

30. Hu C, Li T, Xu Y, Zhang X, Li F, Bai J, et al. CellMarker 2.0: An updated database of manually curated cell markers in human/mouse and web tools based on scRNA-seq data. Nucleic Acids Res [Internet]. 2023 Jan 6 [cited 2025 Apr 24];51(D1):D870–6. Available from: 10.1093/nar/gkac947

31. Andrews TS, Nakib D, Perciani CT, Ma XZ, Liu L, Winter E, et al. Single-cell, single-nucleus, and spatial transcriptomics characterization of the immunological landscape in the healthy and PSC human liver. J Hepatol [Internet]. 2024 May [cited 2024 June 28];80(5):730–43. Available from: https://linkinghub.elsevier.com/retrieve/pii/S0168827824000035

32. Li W, Pan L, Hong W, Ginhoux F, Zhang X, Xiao C, et al. A single-cell pan-cancer analysis to show the variability of tumor-infiltrating myeloid cells in immune checkpoint blockade. Nat Commun [Internet]. 2024 July 21 [cited 2025 Apr 11];15(1):6142. Available from: https://www.nature.com/articles/s41467-024-50478-8

33. Wang AZ, Mashimo BL, Schaettler MO, Sherpa ND, Leavitt LA, Livingstone AJ, et al. Glioblastoma-Infiltrating CD8+ T Cells Are Predominantly a Clonally Expanded GZMK+ Effector Population. Cancer Discov. 2024 June 3;14(6):1106–31.

34. Li K, Zhang R, Wen F, Zhao Y, Meng F, Li Q, et al. Single-cell dissection of the multicellular ecosystem and molecular features underlying microvascular invasion in HCC. Hepatology [Internet]. 2024 June [cited 2025 Feb 12];79(6):1293. Available from: https://journals.lww.com/hep/fulltext/2024/06000/single_cell_dissection_of_the_multicellular.12.aspx

35. Yu Y, Li Y, Zhou L, Cheng X, Gong Z. Hepatic stellate cells promote hepatocellular carcinoma development by regulating histone lactylation: Novel insights from single-cell RNA sequencing and spatial transcriptomics analyses. Cancer Lett [Internet]. 2024 Nov 1 [cited 2025 Feb 12];604:217243. Available from: https://www.sciencedirect.com/science/article/pii/S0304383524006384

36. Zhang DH, Liang C, Hu SY, Huang XY, Yu L, Meng XL, et al. Application of a single-cell-RNA-based biological-inspired graph neural network in diagnosis of primary liver tumors. J Transl Med [Internet]. 2024 Oct 1 [cited 2025 Feb 19];22(1):883. Available from: 10.1186/s12967-024-05670-1

37. Kotliar, D., Curtis, M., Agnew, R. et al. Reproducible single-cell annotation of programs underlying T cell subsets, activation states and functions. Nat Methods 22, 1964–1980 (2025). Available from: 10.1038/s41592-025-02793-1

38. Troulé K, Petryszak R, Cakir B, Cranley J, Harasty A, Prete M, et al. CellPhoneDB v5: Inferring cell-cell communication from single-cell multiomics data. Nat Protoc. 2025 Mar 25;

39. Ziyu Shi, Zhiyuan Song, Heather Steveson-Lerner, Bingning Dong, Haiqing Zhao, LiverDCP: A Disease–Cell–Protein Framework for Multi-scale Modeling of Disease Biology, bioRxiv 2026.08.07.743628; doi: 10.64898/2026.08.07.743628

40. Kim MC, Gate R, Lee DS, Tolopko A, Lu A, Gordon E, Shifrut E, Garcia-Nieto PE, Marson A, Ntranos V, Ye CJ. Method of moments framework for differential expression analysis of single-cell RNA sequencing data. Cell. 2024 Oct 31;187(22):6393–6410.e16. doi: 10.1016/j.cell.2024.09.044. Epub 2024 Oct 24.

41. Ashburner, et al. Gene ontology: tool for the unification of biology. Nat Genet. 2000 May;25(1):25–9. DOI: 10.1038/75556

42. The Gene Ontology Consortium. The Gene Ontology knowledgebase in 2026. Nucleic Acids Res. 2025 Dec 18;gkaf1292. DOI: 10.1093/nar/gkaf1292

43. Gerges, Samar H., et al. “Non-alcoholic fatty liver disease: An overview of risk factors, pathophysiological mechanisms, diagnostic procedures, and therapeutic interventions.” Life Sciences, vol. 271, Apr. 2021, p. 119220, 10.1016/j.lfs.2021.119220.

44. Hong T, Chen Y, Li X, Lu Y. The Role and Mechanism of Oxidative Stress and Nuclear Receptors in the Development of NAFLD. Oxid Med Cell Longev. 2021 Oct 27;2021:6889533. doi: 10.1155/2021/6889533.

45. Martín-Fernández M, Arroyo V, Carnicero C, Sigüenza R, Busta R, Mora N, Antolín B, Tamayo E, Aspichueta P, Carnicero-Frutos I, Gonzalo-Benito H, Aller R. Role of Oxidative Stress and Lipid Peroxidation in the Pathophysiology of NAFLD. Antioxidants (Basel). 2022 Nov 10;11(11):2217. doi: 10.3390/antiox11112217.

46. Arroyave-Ospina JC, Wu Z, Geng Y, Moshage H. Role of Oxidative Stress in the Pathogenesis of Non-Alcoholic Fatty Liver Disease: Implications for Prevention and Therapy. Antioxidants (Basel). 2021 Jan 26;10(2):174. doi: 10.3390/antiox10020174.

47. Rolo AP, Teodoro JS, Palmeira CM. Role of oxidative stress in the pathogenesis of nonalcoholic steatohepatitis. Free Radic Biol Med. 2012 Jan 1;52(1):59–69. doi: 10.1016/j.freeradbiomed.2011.10.003.

48. Loboda A, Damulewicz M, Pyza E, Jozkowicz A, Dulak J. Role of Nrf2/HO-1 system in development, oxidative stress response and diseases: an evolutionarily conserved mechanism. Cell Mol Life Sci. 2016 Sep;73(17):3221–47. doi: 10.1007/s00018-016-2223-0.

49. Akl MG, Li L, Widenmaier SB. Protective Effects of Hepatocyte Stress Defenders, Nrf1 and Nrf2, against MASLD Progression. Int J Mol Sci. 2024 Jul 24;25(15):8046. doi: 10.3390/ijms25158046.

50. Xu D, Xu M, Jeong S, Qian Y, Wu H, Xia Q and Kong X (2019) The Role of Nrf2 in Liver Disease: Novel Molecular Mechanisms and Therapeutic Approaches. Front. Pharmacol. 9:1428. doi: 10.3389/fphar.2018.01428

51. Amir Ajoolabady, Hamid Aslkhodapasandhokmabad, Peter Libby, Jaakko Tuomilehto, Gregory Y.H. Lip, Josef M. Penninger, Des R. Richardson, Daolin Tang, Hao Zhou, Shuyi Wang, Daniel J. Klionsky, Guido Kroemer, Jun Ren,

52. Ferritinophagy and ferroptosis in the management of metabolic diseases, Trends in Endocrinology & Metabolism, Volume 32, Issue 7, 2021, Pages 444–462, ISSN 1043-2760, 10.1016/j.tem.2021.04.010.

53. Hammoutene A, Laouirem S, Albuquerque M, Colnot N, Brzustowski A, Valla D, Provost N, Delerive P, Paradis V; QUID-NASH Research Group. A new NRF2 activator for the treatment of human metabolic dysfunction-associated fatty liver disease. JHEP Rep. 2023 Jul 12;5(10):100845. doi: 10.1016/j.jhepr.2023.100845.

54. Li N, Hao L, Li S, Deng J, Yu F, Zhang J, Nie A, Hu X. The NRF-2/HO-1 Signaling Pathway: A Promising Therapeutic Target for Metabolic Dysfunction-Associated Steatotic Liver Disease. J Inflamm Res. 2024 Nov 3;17:8061–8083

55. Gekakis N, Staknis D, Nguyen HB, Davis FC, Wilsbacher LD, King DP, Takahashi JS, Weitz CJ. Role of the CLOCK protein in the mammalian circadian mechanism. Science. 1998 Jun 5;280(5369):1564–9. doi: 10.1126/science.280.5369.1564

56. Yan J, Wang H, Liu Y, Shao C. Analysis of gene regulatory networks in the mammalian circadian rhythm. PLoS Comput Biol. 2008 Oct;4(10):e1000193. doi: 10.1371/journal.pcbi.1000193. Epub 2008 Oct 10.

57. Lee S, Donehower LA, Herron AJ, Moore DD, Fu L. Disrupting circadian homeostasis of sympathetic signaling promotes tumor development in mice. PLoS One. 2010 Jun 7;5(6):e10995. doi: 10.1371/journal.pone.0010995

58. Padilla J, Osman NM, Bissig-Choisat B, Grimm SL, Qin X, Major AM, Yang L, Lopez-Terrada D, Coarfa C, Li F, Bissig KD, Moore DD, Fu L. Circadian dysfunction induces NAFLD-related human liver cancer in a mouse model. J Hepatol. 2024 Feb;80(2):282–292. doi: 10.1016/j.jhep.2023.10.018. Epub 2023 Oct 27.

59. Xu C, Xu Z, Zhang Y, Evert M, Calvisi DF, Chen X. β-Catenin signaling in hepatocellular carcinoma. J Clin Invest. 2022 Feb 15;132(4):e154515. doi: 10.1172/JCI154515

60. Machado MV. Hepatocellular Carcinoma Around the Clock. Curr Oncol. 2026 Jan 7;33(1):32. doi: 10.3390/curroncol33010032.

61. Mteyrek A, Filipski E, Guettier C, Okyar A, Lévi F. Clock gene Per2 as a controller of liver carcinogenesis. Oncotarget. 2016 Dec 27;7(52):85832–85847. doi: 10.18632/oncotarget.11037

62. M. Qu, G. Zhang, H. Qu, A. Vu, R. Wu, H. Tsukamoto, Z. Jia, W. Huang, H. Lenz, J.N. Rich, & S.A. Kay, Circadian regulator BMAL1::CLOCK promotes cell proliferation in hepatocellular carcinoma by controlling apoptosis and cell cycle, Proc. Natl. Acad. Sci. U.S.A. 120 (2) e2214829120, 10.1073/pnas.2214829120 (2023).

63. Renwen Long, Suoangbaji, Irene Oi-Lin Ng, Daniel Wai-Hung Ho, LiverSCA: A comprehensive and user-friendly cell atlas in human hepatocellular carcinoma, Computational and Structural Biotechnology Journal, Volume 23, 2024, Pages 2740–2745, ISSN 2001-0370, 10.1016/j.csbj.2024.06.031.

64. Li, Z., Zhang, H., Li, Q. et al. GepLiver: an integrative liver expression atlas spanning developmental stages and liver disease phases. Sci Data 10, 376 (2023). 10.1038/s41597-023-02257-1

65. Harpreet Kaur, Sherry Bhalla, Dilraj Kaur, Gajendra PS Raghava, CancerLivER: a database of liver cancer gene expression resources and biomarkers, Database, Volume 2020, 2020, baaa012, 10.1093/database/baaa012

66. Guilliams, Martin, et al. “Spatial Proteogenomics reveals distinct and evolutionarily conserved hepatic macrophage niches.” Cell, vol. 185, no. 2, Jan. 2022, 10.1016/j.cell.2021.12.018.

67. Brancale J, Vilarinho S. A single cell gene expression atlas of 28 human livers Journal of Hepatology. 2021 Jul; 75(1). Available from https://pubmed.ncbi.nlm.nih.gov/34016468/

68. Ma L, Wang L, Khatib SA, Chang CW, Heinrich S, Dominguez DA, Forgues M, Candia J, Hernandez MO, Kelly M, Zhao Y, Tran B, Hernandez JM, Davis JL, Kleiner DE, Wood BJ, Greten TF, Wang XW. Single-cell atlas of tumor cell evolution in response to therapy in hepatocellular carcinoma and intrahepatic cholangiocarcinoma. J Hepatol. 2021 Dec;75(6):1397–1408. doi: 10.1016/j.jhep.2021.06.028.

